# *DEPower*: approximate power analysis with *DESeq2*

**DOI:** 10.64898/2026.02.05.704084

**Authors:** Gennady Gorin, Deek Guruge, Linda Goodman

## Abstract

Rigorous experimental design, including formal power analysis, is a cornerstone of reproducible RNA sequencing (RNA-seq) research. The design of RNA-seq experiments requires computing the minimum sample number required to identify an effect of a particular size at a predefined significance level. Ideally, the statistical test used for the analysis of experimental data should match the test used for sample size determination; however, few tools use the assumptions of the popular differential expression testing framework *DESeq2*, and most opt for simulation-based rather than analytical approaches. Grounded in the *DESeq2* model framework, we derive sample size requirements for both single-cell and bulk RNA-seq experiments delivered as a web-based tool for power analysis, *DEPower*, available at https://poweranalysis-fb.streamlit.app/ that makes rigorous RNA-seq study design accessible to all researchers.

## Analysis

Differential expression (DE) analysis is a ubiquitous part of transcriptomics workflows. In a typical scenario, such an analysis involves constructing a statistical model for the observed gene expression, encoding biologically interesting and incidental covariates, such as the respective effects of a perturbation and the experimental batch, fitting the model, and performing a series of null hypothesis statistical tests to identify genes that display meaningful, high-magnitude differences in expression between the covariates being tested. A wide variety of methods to perform DE testing are available [1–4], but *DESeq2*, a software method originally published in 2014 [5], is particularly widespread and considered a reasonable approach for count-based bulk and single-cell transcriptomics [2, 6], with a robust body of literature interrogating its advantages and caveats [7, 8].

It is often of interest to estimate the experimental sample size necessary to detect a difference of a particular magnitude. In spite of the breadth of studies and software packages that provide relatively sophisticated numerical solutions to this problem [1, 9–16], there are few analytical treatments [10, 17], and we are not aware of any focused specifically on the model used in *DESeq2*. This is somewhat surprising, because the approximate relationship between sample size, effect size, and statistical significance may be derived as an elementary consequence of this model. In this brief manuscript, we fill this lacuna and provide a web app to perform this analysis (Figure 1a). The approach incorporates the *DESeq2* model by constraining the dispersion to a range plausible at the expression level (Figure 1b), producing test statistic (Figure 1c) and *p*-value (Figure 1d) estimates concordant with the full *DESeq2* analysis. By inverting the calculation under this set of assumptions, we identify the corresponding domain of sample sizes.

**Fig 1.**
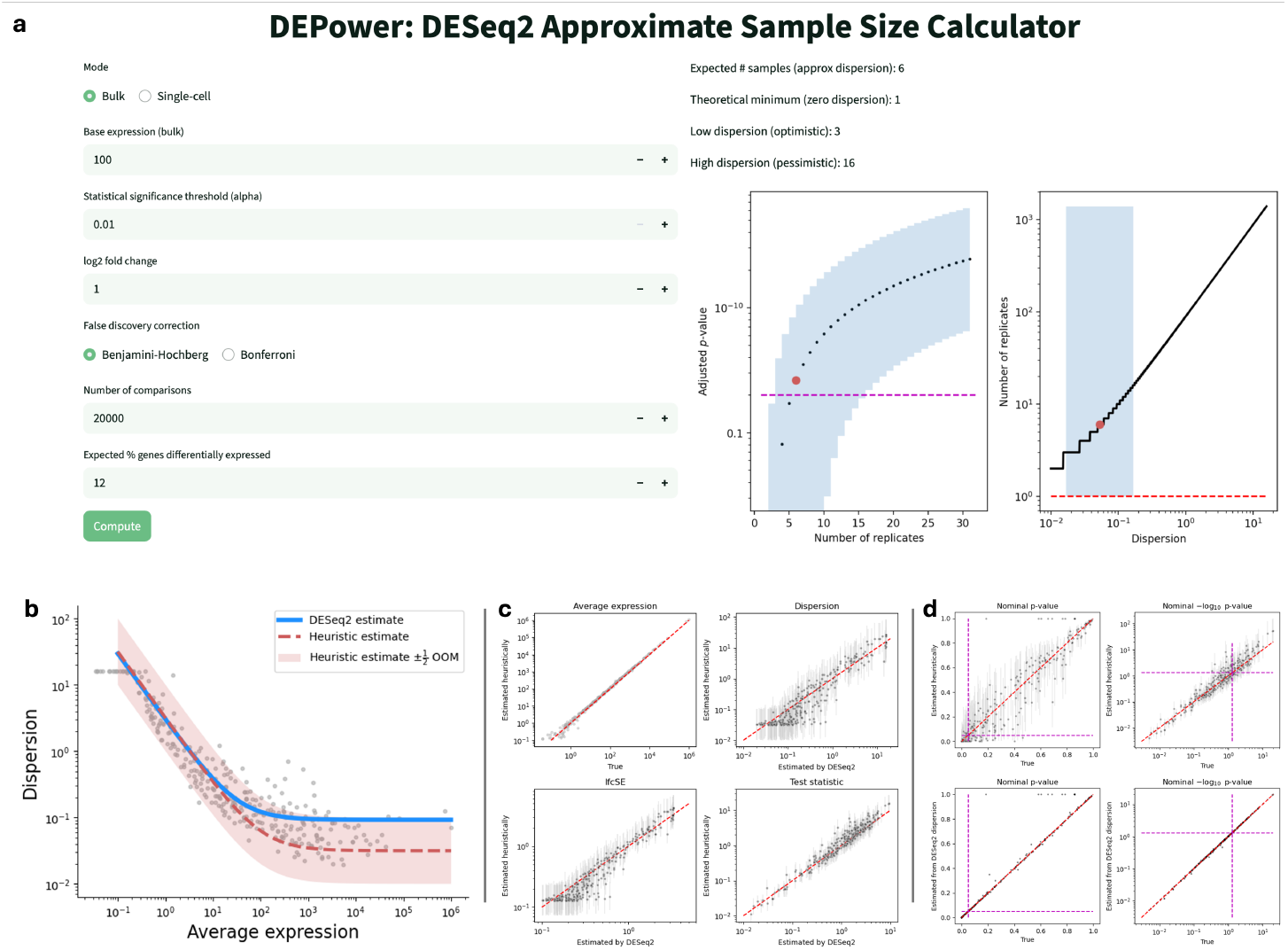
The computation procedure, its implementation, and its comparison to the *DESeq2* procedure. **a**. The web-based calculator interface (left: user-defined inputs; right: computed sample sizes and supporting visualizations). **b**. Dispersion/expression relationship (points: dispersions inferred by *PyDESeq2* ; blue curve: genome-wide dispersion curve inferred by *PyDESeq2* based on data; red line and region: plausible dispersion curve and region estimated by the heuristic procedure). **c**. The relationship between intermediate statistics derived from the heuristic procedure and *PyDESeq2* (points: statistics obtained using the typical-case curve indicated by a red line in **a**; error bars: range of statistics obtained between the upper and lower bounds of the region in **a**; red line: identity). **d**. The relationship between *p*-values derived from the heuristic procedure and *PyDESeq2* (points, top: *p*-values obtained using the typical-case curve indicated by a red line in **a**; error bars, top: range of *p*-values obtained between the upper and lower bounds of the region in **a**; points, bottom: values derived using the *PyDESeq2* dispersion estimate; left: nominal *p*-value, lower is more significant; right: −log_10_ nominal *p*-value, higher is more significant; red line: identity; magenta line: *α* = 0.05).

## Derivation

In the simplest scenario, a *DESeq2* analysis involves a single binary covariate of interest. The nominal *p*-value for a two-sided Wald test of a single parameter under the null hypothesis 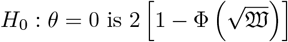, i.e., 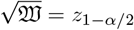 at the two-sided significance level *α*. 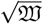, the test statistic, is the square root of the Wald statistic, defined as

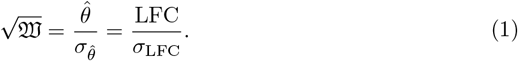

In the context of gene differential expression testing, 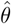 is the estimated log-fold change LFC. As LFC 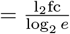, the value is approximately specified by the log_2_ fold change (l_2_fc) one is interested in detecting.

The value *σ*_LFC_ is the standard error of the estimate. For a simple contrast with a balanced design (*n* replicates per condition), and neglecting any variability in read depths between replicates, the square of this quantity is approximately

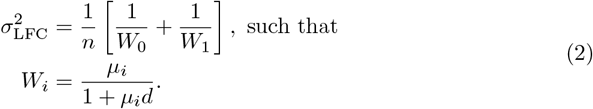

The quantities *W*_0_ and *W*_1_ are per-sample Fisher information matrix contributions to the estimate precision, and are computed by adjusting fitted means *µ*_*i*_ by the gene-specific dispersion *d*. The fitted means are, in turn, obtained directly from the log-fold change, such that *µ*_1_ = *e*^LFC^*µ*_0_.

The theoretical dispersion curve is parametrized according to

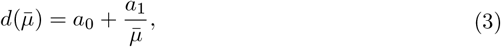

where 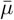 is the average depth-normalized expression. The parameters *a*_0_ and *a*_1_ are obtained from data. It is typical for the parameters to be *d*(1) ≈ *a*_1_ ∈ [1, 10] and 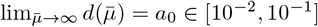. Therefore, a somewhat arbitrary but reasonable choice is the logarithmic midpoint, yielding 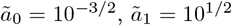. For a given 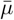, an order-of-magnitude range of dispersions may be obtained by using 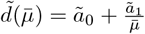, an optimistic low-dispersion case 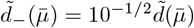, and a pessimistic high-dispersion case 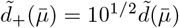.

For the sake of thoroughness, we note that, by default, the dispersion is constrained to the domain [10^−8^, max(*n*, 10)]. The upper limit is generally most relevant in the low-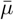 domain. It is straightforward to solve the corresponding equation,

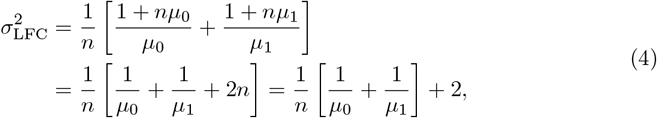

which only provides a real-valued solution for 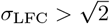. In other words, if 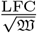 is below this threshold, the desired significance threshold cannot be satisfied at any finite sample size. However, this edge case only becomes relevant at *d*_+_ = 10 or 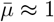, a regime more appropriate for targeted methods, such as quantitative PCR.

It remains to estimate 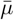, which is the average of normalized counts across all samples. Once again neglecting systematic depth variability, this quantity computed as

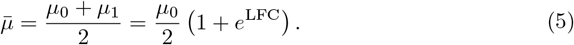

In the unbalanced case with *n*_0_ and *n*_1_ replicates per condition, the first line of Equation 2 reads

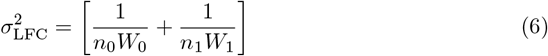

and Equation 5 reads

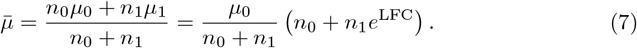

In other words, to estimate the sample size required to achieve a particular nominal *p*-value and log-fold change, one may

1. Find the requisite test statistic 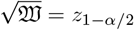 for the *α* of interest.
2. Assume a particular log-fold change and solve for the standard error using Equation 1.
3. Assume a particular base expression level *µ*_0_ and use Equation 5 to solve for the overall average 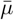.
4. Estimate the dispersion *d* based on Equation 3, optionally considering a domain of dispersions.
5. Obtain weights *W* from the second line of Equation 2.
6. Finally, use the first line of Equation 2 to solve for the necessary sample size *n*.

In a genome-wide assay, one is typically interested in detecting expression changes in multiple genes, requiring false discovery control. In the most conservative case, the Bonferroni correction can be implemented by substituting the *p* of interest by 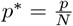, where *N* is the total number of comparisons. However, *DESeq2* natively uses the Benjamini–Hochberg procedure for false discovery rate control, such that an *adjusted p*-value, 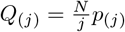, is compared against the predefined significance level *α*, where *j* = 1, …, *N* denotes the rank of the *p*-value. This quantity cannot be estimated, as it is dependent on the data, but the procedure may nevertheless be followed by using a value 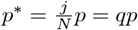, i.e., assuming the *p*-value is at quantile *q* with respect to the rest of the *p*-value distribution, where *q* may be loosely approximated by a hypothetical fraction of true non-null genes [18].

This approach is sufficient for order-of-magnitude, back-of-the-envelope calculations. For instance, if one is interested in designing a single-nucleus sequencing experiment to investigate relatively high-expression genes (e.g., average expression *>* 0.1 molecules per cell) in a rare cell type (e.g., 1% of cells in an experiment), the expected pseudobulk expression in a 10,000-cell sample is 10. Identifying a log_2_ fold change of 1 at a *nominal p*-value threshold of 0.05 in this scenario would require 6 samples per condition (3/14 for the low-/high-dispersion cases). Assuming only some 10,000 genes are expressed highly enough to analyze, achieving this level under the Bonferroni correction would require 28 samples (14/74). Finally, under the Benjamini–Hochberg procedure, coarsely assuming *q* = 0.1 for the genes under consideration, the experiment would require 22 (11/58) samples. In other words, even under the most optimistic assumptions of this idealized scenario, achieving the desired level of statistical significance would require dozens of samples. These study sizes may be unachievable or cost-prohibitive, implying that standard “unbiased” single-cell RNA sequencing is insufficient to investigate this cell population at the specified significance, and cell type enrichment is mandatory.

The accuracy of the heuristic procedure relies on the validity of its assumptions. The neglect of read depth differences between conditions is an oversimplification, but typically a minor one. On the other hand, the *ad hoc* dispersion estimation tends to incur more severe errors. In Figure 1b-d, we illustrate its performance on the “forward” estimation problem on a small dataset quantifying the abundance of microRNAs in the livers of thirteen-lined ground squirrels during winter torpor and summer [19], using *PyDESeq2*, a Python re-implementation of *DESeq2*, as a baseline [20]. The heuristic dispersion estimate generally agrees with the trend inferred using the *DESeq2* process, although individual genes may lie outside these bounds (Figure 1b). The uncertainty of the estimate can be propagated to the computed statistics (Figure 1c), including the *p*-values (Figure 1d, top), which are broadly consistent with the full procedure. Using the “true” dispersion estimate from *PyDESeq2* instead of the heuristic estimate, while retaining the rest of the procedure, produces *p*-value estimates that are nearly identical to those obtained by the full procedure (Figure 1d, bottom). As discussed elsewhere, one can avoid this procedure if pilot data are available [12, 15], as such data can provide more relevant estimates of the expression levels and the dispersion distribution.

The assumptions specific to the approximation are compounded with the usual assumptions of null hypothesis statistical testing under the *DESeq2* framework. In particular, the Wald test relies on the asymptotic convergence of the test statistic to the chi-square distribution, which is generally a reasonable assumption for moderate-to-large sample sizes, but may be less reliable for smaller sample sizes. In addition, any use of *DESeq2* implicitly assumes that the negative binomial statistical model is appropriate under the given comparison; if the model is violated, additional covariates (such as distinct batches) are present, or outliers have been produced due to technical effects, the effective sample size may need to be higher than predicted. Therefore, the results of the sample size analysis should be seen as a lower bound under the ideal-case scenario.

## Conclusion

In this report, we provide a simple strategy for estimating sample sizes necessary to achieve a particular level of statistical power in RNA sequencing experiments, presupposing that the planned downstream analysis uses the *DESeq2* statistical model and that the variability in data is within typical bounds. The procedure outlined here may be adapted to specific circumstances based on the range of available experimental data, or followed directly under the outlined assumptions.

### Calculator

To facilitate experiment design, we provide the *DEPower* calculator that implements the strategy at https://poweranalysis-fb.streamlit.app/, shown in Figure 1a. To compute the sample size, the user needs to specify the experiment design (bulk or single-cell) and the variables; for bulk RNA sequencing, these include the base expression *µ*_0_, statistical significance threshold *α*, log-fold change l_2_fc, and the false discovery strategy. The calculator reports the sample sizes required to achieve the desired level of significance using the order-of-magnitude estimates for dispersion outlined above. Additionally, to clarify the method assumptions, it visualizes the dependence of the minimum achievable *p*-value on the sample size and the dependence of *n* on the dispersion. The open-source calculator is implemented and deployed using Streamlit.

## Methods

To demonstrate the performance of the approximate method, we compare the statistics derived from this procedure to those obtained by the usual *DESeq2* workflow, implemented using *PyDESeq2* 0.4.8 [20]. In this largely qualitative illustration, we attempt to characterize the effects of the simplifications on the results, using the “forward” procedure — i.e., performing a statistical test using only the portion of the available information relevant to experiment design — as a simple baseline that can be immediately and intuitively compared to the results of the full workflow. In this scenario, *µ*_0_ is the average of raw counts in the summer condition and l_2_fc is somewhat arbitrarily set to the log_2_ fold change inferred by *PyDESeq*2.

We obtained the read counts for mitochondrial microRNA profiles from the Robichaud et al. study [19] from the Gene Expression Omnibus series GSE254223 and filtered them for summer and torpid (TSL and TTL) thirteen-lined ground squirrels, with eight animals disclosed per group. We used the default parameters to fit the data and perform statistical testing.

Figure 1c compares the intermediate statistics obtained from the heuristic procedure to those obtained by *PyDESeq2*. In the top left panel, we compare 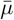 obtained from Equation 5 to that obtained by directly computing the average of normalized counts (baseMean). In the top right panel, we compare the heuristic dispersion estimates (red line and region in Figure 1b) to the ones obtained by the full inference procedure (points in Figure 1b, obtained by shrinkage toward the best-fit line indicated in blue). These estimates are propagated to the standard error of the log-fold change, in base 2 units, using Equation 2, as shown in the bottom left panel, as well as the test statistic, using Equation 1, as shown in the bottom right panel. The test statistic can be transformed into an estimate of nominal *p*-value, as shown in the top panels of Figure 1d. As may be expected, the deviations from the *p*-value obtained through the full procedure are overwhelmingly explained by the *ad hoc* dispersion estimate: if the *PyDESeq2* estimate is used, the results agree with the full procedure.

### An aside on power analysis as a detector of scientific fraud

In addition to *de novo* design, this approach may be suitable for detecting research fraud. Above, we discuss constraining sample sizes for a proposed experiment given a particular *p*-value and effect size. Using the same mathematics, it is possible to scrutinize published experiments and check whether extreme results — for instance, a *p*-value of 10^−200^ at a log_2_ fold change of 1 based on *n* = 3 samples — are mathematically plausible, and provide supporting evidence for potential cases of analysis errors or misconduct if not. However, this approach is unlikely to be broadly fruitful for this use case, for the following reasons:

- The approach is considerably more challenging and requires more manual work than the detection of other forms of scientific misconduct visible through duplicated images, “tortured phrases” intended to bypass plagiarism detection, irrelevant citations, or hallmarks of non-human writing [21–24].
- Inconsistencies are suggestive, but not conclusive; for a gene with sufficiently high expression and low dispersion, one could, in principle, achieve arbitrarily low *p*-values, in contrast to, e.g., the impossibility of achieving a mean of 1.25 by averaging two integers [25, 26].
- Intentional misconduct is not readily distinguishable from misuse of the statistical tool, such as normalizing, transforming, and rounding data without adapting the statistical test appropriately.
- Low-*n* analyses, where these constraints are relevant, are more frequent in studies reporting and analyzing newly generated data; on the other hand, fraudulent studies more frequently draw on large public datasets, such as The Cancer Genome Atlas, with dozens or hundreds of replicates [27, 28].
- In the single-cell sequencing field, the violation of statistical constraints is typically the result of the improper interpretation of cells derived from the same donor organism as independent biological replicates, or pseudoreplication, frequently exacerbated by the lack of multiple hypothesis correction [29, 30].
- The falsification of statistical test results is considerably more time-consuming, and thus less plausible, than misrepresentation of the data used for the test. In addition, data may be fabricated wholesale [31].

Given this context and the associated constraints, it does not appear that this approach can be applied to the rapidly growing problem of research misconduct [23] at the appropriate scale and level of confidence. Nevertheless, it may be of some value for performing diligence checks on studies with far-reaching claims based on a small number of samples, and implausible *p*-values may be an indicator that other statistical procedures warrant scrutiny.

## Data and code availability

A web app that implements the calculations outlined in this report is available at https://poweranalysis-fb.streamlit.app/. A notebook that reproduces the results shown in Figure 1, as well as the source code of the web app, are available at https://github.com/Fauna-Bio/GGG_2026.

## Acknowledgments

L.G. is a co-founder and CTO of Fauna Bio. G.G. and D.G. are employees of Fauna Bio. The authors would like to acknowledge Reese Richardson, Evelyn Tran, and Hanbin Lee for their assistance in the preparation of this manuscript and tool.

## References

1. Schurch NJ, Schofield P, Gierliński M, Cole C, Sherstnev A, Singh V, et al. How many biological replicates are needed in an RNA-seq experiment and which differential expression tool should you use? RNA. 2016;22(6):839–851. doi:10.1261/rna.053959.115.

2. Baik B, Yoon S, Nam D. Benchmarking RNA-seq differential expression analysis methods using spike-in and simulation data. PLOS ONE. 2020;15(4):e0232271. doi:10.1371/journal.pone.0232271.

3. Tang M, Sun J, Shimizu K, Kadota K. Evaluation of methods for differential expression analysis on multi-group RNA-seq count data. BMC Bioinformatics. 2015;16(1):360. doi:10.1186/s12859-015-0794-7.

4. Zhang ZH, Jhaveri DJ, Marshall VM, Bauer DC, Edson J, Narayanan RK, et al. A Comparative Study of Techniques for Differential Expression Analysis on RNA-Seq Data. PLOS ONE. 2014;9(8):e103207. doi:10.1371/journal.pone.0103207.

5. Love MI, Huber W, Anders S. Moderated estimation of fold change and dispersion for RNA-seq data with DESeq2. Genome Biology. 2014;15(12):550. doi:10.1186/s13059-014-0550-8.

6. Luecken MD, Theis FJ. Current best practices in single-cell RNA-seq analysis: a tutorial. Molecular Systems Biology. 2019;15(6):e8746. doi:10.15252/msb.20188746.

7. Lee H, Han B. Pseudobulk with proper offsets has the same statistical properties as generalized linear mixed models in single-cell case-control studies. Bioinformatics. 2024;40(8):btae498. doi:10.1093/bioinformatics/btae498.

8. Squair JW, Gautier M, Kathe C, Anderson MA, James ND, Hutson TH, et al. Confronting false discoveries in single-cell differential expression. Nature Communications. 2021;12(1):5692. doi:10.1038/s41467-021-25960-2.

9. Ching T, Huang S, Garmire LX. Power analysis and sample size estimation for RNA-Seq differential expression. RNA. 2014;20(11):1684–1696. doi:10.1261/rna.046011.114.

10. Bi R, Liu P. Sample size calculation while controlling false discovery rate for differential expression analysis with RNA-sequencing experiments. BMC Bioinformatics. 2016;17(1):146. doi:10.1186/s12859-016-0994-9.

11. Li CI, Shyr Y. Sample size calculation based on generalized linear models for differential expression analysis in RNA-seq data. Statistical Applications in Genetics and Molecular Biology. 2016;15(6):491–505. doi:10.1515/sagmb-2016-0008.

12. Poplawski A, Binder H. Feasibility of sample size calculation for RNA-seq studies. Briefings in Bioinformatics. 2018;19(4):713–720. doi:10.1093/bib/bbw144.

13. Li WV, Li JJ. A statistical simulator scDesign for rational scRNA-seq experimental design. Bioinformatics. 2019;35(14):i41–i50. doi:10.1093/bioinformatics/btz321.

14. Su K, Wu Z, Wu H. Simulation, power evaluation and sample size recommendation for single-cell RNA-seq. Bioinformatics. 2020;36(19):4860–4868. doi:10.1093/bioinformatics/btaa607.

15. Schmid KT, Höllbacher B, Cruceanu C, Böttcher A, Lickert H, Binder EB, et al. scPower accelerates and optimizes the design of multi-sample single cell transcriptomic studies. Nature Communications. 2021;12(1):6625. doi:10.1038/s41467-021-26779-7.

16. Jeon H, Xie J, Jeon Y, Jung KJ, Gupta A, Chang W, et al. Statistical Power Analysis for Designing Bulk, Single-Cell, and Spatial Transcriptomics Experiments: Review, Tutorial, and Perspectives. Biomolecules. 2023;13(2):221. doi:10.3390/biom13020221.

17. Hart SN, Therneau TM, Zhang Y, Poland GA, Kocher JP. Calculating Sample Size Estimates for RNA Sequencing Data. Journal of Computational Biology. 2013;20(12):970–978. doi:10.1089/cmb.2012.0283.

18. Benjamini Y, Hochberg Y. Controlling the False Discovery Rate: A Practical and Powerful Approach to Multiple Testing. Journal of the Royal Statistical Society Series B (Methodological). 1995;57(1):289–300.

19. Robichaud K, Duffy B, Staples JF, Craig PM. Mitochondrial microRNA profiles are altered in thirteen-lined ground squirrels (Ictidomys tridecemlineatus) during hibernation. Physiological Genomics. 2024;56(8):555–566. doi:10.1152/physiolgenomics.00017.2024.

20. Muzellec B, Teleńczuk M, Cabeli V, Andreux M. PyDESeq2: a python package for bulk RNA-seq differential expression analysis. Bioinformatics. 2023;39(9):btad547. doi:10.1093/bioinformatics/btad547.

21. Else H. ‘Tortured phrases’ give away fabricated research papers. Nature. 2021;596(7872):328–329. doi:10.1038/d41586-021-02134-0.

22. Bik EM, Casadevall A, Fang FC. The Prevalence of Inappropriate Image Duplication in Biomedical Research Publications. mBio. 2016;7(3):e00809–16. doi:10.1128/mBio.00809-16.

23. Richardson RAK, Hong SS, Byrne JA, Stoeger T, Amaral LAN. The entities enabling scientific fraud at scale are large, resilient, and growing rapidly. Proceedings of the National Academy of Sciences. 2025;122(32):e2420092122. doi:10.1073/pnas.2420092122.

24. Liang W, Zhang Y, Wu Z, Lepp H, Ji W, Zhao X, et al. Mapping the Increasing Use of LLMs in Scientific Papers; 2024. Available from: http://arxiv.org/abs/2404.01268.

25. Brown NJL, Heathers JAJ. The GRIM Test: A Simple Technique Detects Numerous Anomalies in the Reporting of Results in Psychology. Social Psychological and Personality Science. 2017;8(4):363–369. doi:10.1177/1948550616673876.

26. Crone G. Can statistical methods reliably detect fraudulent data? Examining the utility of p-value analyses, extreme effect sizes, GRIM, and GRIMMER. York University. Toronto, Ontario; 2025. Available from: https://hdl.handle.net/10315/43052.

27. Spick M, Onoja A, Harrison C, Stender S, Byrne J, Geifman N. Quantifying new threats to health and biomedical literature integrity from rapidly scaled publications and problematic research; 2025. Available from: https://www.medrxiv.org/content/10.1101/2025.07.07.25331008v1.

28. Suchak T, Aliu AE, Harrison C, Zwiggelaar R, Geifman N, Spick M. Explosion of formulaic research articles, including inappropriate study designs and false discoveries, based on the NHANES US national health database. PLOS Biology. 2025;23(5):e3003152. doi:10.1371/journal.pbio.3003152.

29. Zimmerman KD, Espeland MA, Langefeld CD. A practical solution to pseudoreplication bias in single-cell studies. Nature Communications. 2021;12(1):738. doi:10.1038/s41467-021-21038-1.

30. Mukamel EA, Yu Z. False positives in study of memory-related gene expression. Nature. 2025;642(8066):E1–E3. doi:10.1038/s41586-025-08988-y.

31. Bradshaw MS, Payne SH. Detecting fabrication in large-scale molecular omics data. PLOS ONE. 2021;16(11):e0260395. doi:10.1371/journal.pone.0260395.

